# Bioinformatic analysis and experimental validation of ursolic acid’s effects on *Staphylococcus aureus*-induced osteomyelitis

**DOI:** 10.1101/2025.10.10.681659

**Authors:** Chengshuo Deng, Shuangli Chen, Wenen Huang, Longjie Miao, Yun Zhao, Huijun Zhu, Yumei Zhou, Lijun Zhao, Helu Liu

## Abstract

Osteomyelitis (OM), particularly methicillin-resistant *Staphylococcus aureus* (MRSA)-induced OM, remains a serious clinical challenge due to complex pathogenesis and rising antibiotic resistance. This study aimed to elucidate the therapeutic mechanism of ursolic acid (UA) against MRSA-induced OM using integrated network pharmacology, bioinformatics, and experimental validation. Through target prediction and multi-database mining, we identified 57 common targets of UA and OM. ML algorithms, including Least Absolute Shrinkage and Selection Operator (LASSO) and random forest, pinpointed three core genes: Neutrophil Elastase (*ELANE*), Lactoferrin (*LTF*), and *S100A12*. Enrichment analyses revealed significant involvement in neutrophil extracellular trap (NET) formation, inflammatory response, and immune regulation. Molecular docking and dynamics simulations confirmed stable binding between UA and *ELANE*. *In vitro*, UA exhibited antibacterial activity against clinical MRSA isolates (MIC = 16–64 μg/mL) and showed synergistic effects with Penicillin (FICI = 0.125–0.5). These results demonstrate that UA combats OM through immunomodulation, anti-inflammatory actions, and direct antibacterial activity, providing a foundation for its use as an adjuvant therapy against MRSA-related OM.

## Introduction

Osteomyelitis (OM), an inflammatory condition of bone tissue, is primarily triggered by the invasion of bacterial pathogens into the skeletal system. Bacterial OM poses particular therapeutic challenges, partly due to the extensive antibiotic resistance of *Staphylococcus aureus* (*S. aureus*)— a predominant Gram-positive pathogen (1). The disease triggers pathological bone remodeling, leading to the isolation of infectious foci from innate immune effector cells and systemic antibiotics. Thus, clinical management often requires prolonged antibiotic regimens combined with surgical debridement of infected and necrotic tissue (2). Even with these aggressive measures, many patients develop chronic infections or associated complications. A deeper understanding of the mechanisms through which bacteria invade bone, survive within bone tissue, and induce pathological bone remodeling will facilitate the development of novel strategies for preventing and treating OM, as well as mitigating its complications (3).

Based on disease duration, OM is generally classified as acute or chronic; based on the route of infection, it may be categorized as hematogenous or contiguous. During bone infection, serum levels of various cytokines [such as interleukin-6 (IL-6), IL-8, interleukin-1β (IL- 1β), and IL-12 (p70)], angiogenic factors (e.g., vascular endothelial growth factor, VEGF), and acute-phase proteins (e.g., C-reactive protein, CRP) are elevated, which holds clinical significance for diagnosis and treatment.

However, there remains a need to identify specific infection biomarkers with diagnostic value (4).

According to the mechanism of infection, OM can be further divided into hematogenous and non-hematogenous types. In hematogenous OM, bacteria are seeded in the bone secondary to bloodstream infection; this form is most common in children, the elderly, and immunocompromised individuals (5). Non-hematogenous OM results from direct inoculation following surgery or trauma, or from the spread of adjacent soft tissue and joint infections. Methicillin-susceptible *Staphylococcus aureus* (MSSA) is the most frequent pathogen in all types of OM, followed by *Pseudomonas aeruginosa* and methicillin-resistant *Staphylococcus aureus* (MRSA). Hematogenous OM is typically monomicrobial and may involve aerobic Gram-negative bacilli, while injection drug users are susceptible to infections with *P. aeruginosa* or *Serratia marcescens*. In vertebral OM— the most common form of hematogenous OM—5%–10% of cases are polymicrobial. Blood cultures may be negative if OM develops after clearance of bacteria from the bloodstream. Non-hematogenous OM can be polymicrobial; besides *S. aureus*, coagulase-negative staphylococci, aerobic Gram-negative bacteria, and anaerobes are also major pathogens. Polymicrobial infections in diabetic foot infections and pressure ulcer infections may include *Streptococcus* and *Enterococcus* species. Moreover, antimicrobial resistance remains a challenge in OM treatment, with up to 50% of *S. aureus* OM cases caused by MRSA strains (6).

*S. aureus* possesses a wide array of virulence factors that facilitate invasion and survival within host tissues (7–9). These include adhesins, cytolytic toxins and exoenzymes, immune evasion factors, and superantigens. Particularly important in the pathogenesis of OM are microbial surface components recognizing adhesive matrix molecules (MSCRAMMs)—a group of adhesins with IgG-like subdomains capable of binding extracellular matrix components such as collagen, fibrinogen, bone sialoprotein, and fibronectin. The staphylococcal adhesin Cna, which binds collagen, contributes to the pathogenesis of OM (10, 11), although clinical isolates lacking collagen-binding ability have been recovered from human OM cases (12).

A well-studied key aspect of *S. aureus* pathogenesis in OM is biofilm formation, which has been systematically reviewed (13, 14). *S. aureus*, the primary pathogen in OM, invades bone through various mechanisms. It can enter bone via hematogenous spread or direct contamination (e.g., after fracture or surgery). Surface adhesins such as fibronectin-binding proteins A and B (FnBPA/B), collagen adhesin (Cna), and staphylococcal protein A (SpA) (15) enable adhesion to and invasion of osteoblasts by binding host receptors or extracellular matrix proteins. Upon invasion, *S. aureus* activates multiple signaling pathways, including EGFR/FAK and c-Src (16), promoting bacterial uptake and intracellular proliferation while disrupting normal osteoblast function—leading to reduced proliferation, increased apoptosis, and impaired mineralization. Furthermore, *S. aureus* infection induces osteoblasts to secrete various inflammatory mediators and cytokines, such as IL-6, IL-12, and RANKL (17), which enhance immune cell recruitment and activation and promote osteoclast formation and activity, resulting in bone resorption. The ability of *S. aureus* to form biofilms further reduces osteoblast activity and increases RANKL production, exacerbating bone loss (18). Additionally, *S. aureus* can persist intracellularly as small colony variants (SCVs), which exhibit enhanced intracellular persistence and antibiotic resistance, enabling long-term survival within host cells and immune evasion, thereby contributing to chronic OM (19, 20). In summary, *S. aureus* promotes the development and progression of OM through direct invasion of bone cells, activation of inflammatory responses, and formation of biofilms and SCVs.

Ursolic acid (UA) exhibits a spectrum of pharmacological activities centered around antioxidant, anti-inflammatory, and anticancer effects, with additional hepatoprotective, hypoglycemic, antiviral, antibacterial, and antifibrotic properties. Although not yet approved for clinical use, UA is in preclinical and early clinical development, and a topical formulation has been approved in Japan for skin cancer prevention. Researchers have developed a nanostructured lipid carrier loaded with *Olea europaea* leaf extract to improve the transdermal delivery of UA for anti-arthritis therapy (21). Previous studies have shown that UA inhibits osteoclast formation: under induction by M-CSF and RANKL, bone marrow monocytes differentiate into mature osteoclasts (TRAP-positive multinucleated cells). Treatment with 5 μM UA significantly reduced the number of multinucleated cells and the area of TRAP-positive staining. Transfection with IKKβC179A restored osteoclastogenesis, indicating that UA suppresses osteoclast differentiation via IKKβ Cys-179.

Machine learning (ML) employs algorithms to generate predictions from input data, offering objectivity and accuracy surpassing manual computation. Network pharmacology, grounded in systems biology and pharmacology, utilizes techniques such as genomics and network analysis to explore complex "drug–gene–target–disease" interactions. By integrating multidimensional approaches to understand the molecular basis of diseases and predict pharmacological mechanisms, network pharmacology enables the evaluation of drug efficacy, mechanisms of action, and potential adverse effects.

Research on OM faces challenges due to its reliance on traditional principles of Chinese medicine pharmacology. However, the application of network pharmacology offers a feasible approach to investigate pharmacological mechanisms using big data and computational techniques (22, 23). In previous work, we utilized network pharmacology to identify Neutrophil Elastase (*ELANE*) as a target of UA. By examining nodes and pathways within biological networks, we analyzed network characteristics and elucidated the drug’s mechanism of action, providing an effective methodology for Traditional Chinese Medicine (TCM) development. Network pharmacology is widely employed in TCM research, including the screening of active compounds and potential targets, unraveling the mechanisms of complex herbal prescriptions, treating diseases, and exploring the properties of multi-target agents. The integration of TCM databases with computational software has enhanced the level of TCM research and contributed significantly to its internationalization.

Here, we applied network pharmacology to investigate potential therapeutic targets against OM. We subsequently analyzed PPI among the Kyoto Encyclopedia of Genes and Genomes (KEGG) pathways and target clusters, performed Gene Ontology (GO) functional and pathway enrichment analyses, and examined correlations between potential targets and disease- and aging-related genes. Additionally, we employed LASSO regression and Random Forest (RF) algorithms to construct a target screening model. Core targets were further validated using molecular docking and molecular dynamics (MD) simulations. This comprehensive study aims to clarify the complete molecular mechanism underlying the action against *S. aureus* infection-induced OM. We also evaluated the clinical significance of the core targets using datasets from the Gene Expression Omnibus (GEO) database. The analysis workflow of this study is illustrated in the Fig 1 (24).

**Fig 1.**
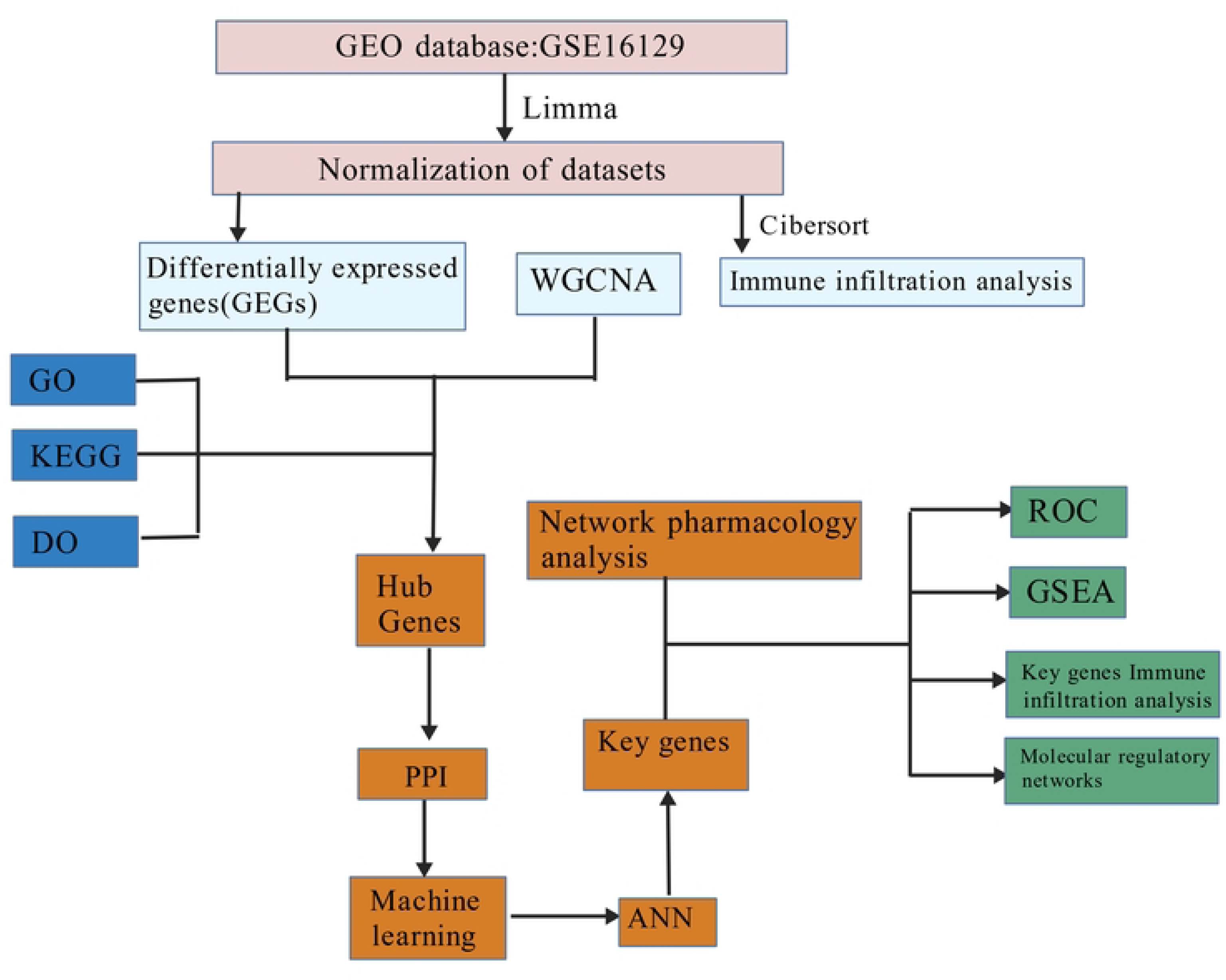
The analysis workflow of this study.

## Materials and methods

### Screening of bioactive compound targets and disease-related targets

This study integrated multiple database resources to predict the potential targets of UA. First, the Simplified Molecular Input Line Entry System (SMILES) sequence and the two-dimensional structural (SDF) file of UA were retrieved from the PubChem database (https://pubchem.ncbi.nlm.nih.gov/) (25). Subsequently, a dual-platform strategy was employed for target prediction: (1) The SMILES sequence was submitted to the SwissTargetPrediction database (http://www.swisstargetprediction.ch/) (26), and potential targets with a probability value greater than 0 were selected. (2) The SDF file was submitted to the PharmMapper online platform (https://www.lilab-ecust.cn/pharmmapper/) (27) for pharmacophore matching and target identification. All targets obtained from both platforms were converted and standardized using the UniProt database (https://www.uniprot.org/) (28). The union of these targets constituted the final set of predicted UA targets.

Simultaneously, using the search terms "osteomyelitis" and "*Staphylococcus aureus* infection", a systematic retrieval was performed across the GeneCards (https://www.genecards.org/) (29), OMIM (https://www.omim.org/) (30), and DisGeNET (https://www.disgenet.org/) (31) databases to collect disease-related target genes. To elucidate the therapeutic relevance of UA in *S. aureus*-infectious OM, the VennDiagram package (v1.7.3) in R (v4.4.1) was used to generate Venn diagrams illustrating the intersections among UA targets, OM-related targets, and *S. aureus* infection-related targets.

### Construction of Protein-Protein Interaction (PPI) network

The intersection targets of UA and the disease identified in Section 2.1 were imported into the STRING database (https://string-db.org/) (32), with the species set as "Homo sapiens" and a confidence threshold > 0.4, to construct a PPI network. The resulting network data were then imported into Cytoscape software (v3.8.0) for visualization and further analysis (33). To identify key hub genes within the network, the CytoHubba plugin in Cytoscape was utilized (34). The importance of each node was evaluated based on its topological properties, primarily using three parameters: Degree (number of edges directly connected to the node), Betweenness Centrality (the proportion of all shortest paths passing through the node), and Closeness Centrality (the reciprocal of the average shortest path length from the node to all other nodes in the network). Based on these metrics, the top-ranked hub genes were selected as potential key targets for subsequent in-depth analysis.

### GO and KEGG pathway enrichment analysis

GO provides a structured framework for functional annotation in bioinformatics, systematically interpreting gene functions from three aspects: Molecular Function (MF), Biological Process (BP), and Cellular Component (CC). The KEGG complements GO by mapping gene functions onto molecular networks through its PATHWAY database, visualizing key metabolic and regulatory pathways.

In this study, the DAVID database (https://david.ncifcrf.gov/) was utilized to perform GO and KEGG pathway enrichment analyses on the common targets associated with OM and *S. aureus* infection (35). The identifier type was set to "OFFICIAL_GENE_SYMBOL" and the species to "Homo sapiens", with a significance threshold of *P* < 0.05. The enrichment results were visualized using the bioinformatics online platform (http://www.bioinformatics.com.cn/), generating statistical plots for BP, CC, MF, and KEGG pathways.

To further clarify the regulatory relationships between targets and pathways, the target gene list was submitted to the UniProt database (https://www.uniprot.org) to convert gene symbols into UniProtKB accession numbers, from which corresponding KEGG identifiers were retrieved. Using the "Color" function in the KEGG Mapper online tool (https://www.genome.jp/kegg/mapper/), the targets were mapped and color-highlighted in relevant pathway diagrams to visually illustrate their potential functional sites in pathways related to *S. aureus*-infectious OM, thereby providing a basis for subsequent mechanistic analysis.

### Correlation analysis of core target genes and OM- related genes

Gene expression profile data related to OM were obtained from the GEO (https://www.ncbi.nlm.nih.gov/geo/) database (36). The data retrieval strategy was as follows: (1) species restricted to Homo sapiens; (2) experimental data type limited to microarray; (3) inclusion of paired samples from both OM patients and healthy controls; and (4) a total sample size of no fewer than 10 (with at least 5 per group). After systematic screening, which excluded datasets from animal model studies, single- group designs, and non-microarray data types, dataset GSE16129 (37) was ultimately included as this study’s OM gene expression dataset.

### Data preprocessing and screening of differentially expressed genes (DEGs)

This study performed gene expression analysis based on the GSE161284 dataset (Platform: GPL96). First, the gene expression matrix was extracted from this dataset, and gene probes were annotated and matched with expression values using the corresponding platform annotation file. Differential expression analysis was conducted using the limma package (38) to identify DEGs between the group of patients with *S. aureus*-induced OM and the healthy control group. The criteria for screening DEGs were set as an adjusted *P*-value < 0.05 and an absolute log2 fold change (|log2FC|) > 0.5. The screening results were visualized using a volcano plot. Furthermore, the top 30 most significantly up- regulated and top 30 most significantly down-regulated DEGs were selected for inter-sample correlation analysis, and a heatmap was generated to display their expression patterns visually.

### Weighted Gene Co-expression Network Analysis (WGCNA)

This study employed the "WGCNA" package in R to construct a weighted gene co-expression network, aiming to identify co-expression modules with significant biological relevance and to explore their associations with disease phenotypes. The specific analytical workflow was as follows: First, genes with the top 50% variance in expression levels were selected for subsequent network construction based on the degree of gene expression variability. Second, the pickSoftThreshold function was used to calculate the optimal soft-thresholding power β within the range of 1 to 20 to approximate a scale-free network topology, with the scale-free topology fitting index R² threshold set to 0.9. Subsequently, the adjacency matrix between genes was transformed into a topological overlap matrix (TOM), and the corresponding dissimilarity measure (1-TOM) was computed. Then, hierarchical clustering combined with a dynamic tree-cutting algorithm was applied to identify co-expression modules, with the minimum module size set to 50 genes and a module merging threshold set at 0.25. Finally, gene significance (GS) and module membership (MM) were calculated to evaluate the correlation between each module and disease status. Genes most significantly associated with the disease within the modules were extracted for further analysis.

### Functional enrichment analysis of intersecting genes

This study performed systematic functional annotation using the "clusterProfiler" R package (39), including GO enrichment analysis and KEGG pathway enrichment analysis to elucidate the biological functions of the DEGs associated with OM. The GO analysis covered three categories: BP, CC, and MF. A statistical significance threshold of *P* < 0.05 was applied to identify key BP, cellular structures, MF, and signal transduction pathways significantly associated with the DEGs.

### Machine learning

To identify core candidate genes for OM, this study employed a multi-algorithm integration strategy. First, feature selection was performed using LASSO regularization regression implemented via the R package "glmnet" (40), while the "randomForest" package (41) was used to run the RF algorithm to assess variable importance. Subsequently, the intersection of results from both algorithms was identified, and Venn diagrams were employed to visualize overlapping genes. Genes commonly selected by both methods were identified as candidate biomarkers for OM. To validate the reliability of these candidate genes, their expression distribution differences in the training dataset were visualized using box plots. Additionally, the "pROC" package (42) was utilized to generate receiver operating characteristic (ROC) curves and calculate the area under the curve (AUC) to evaluate the diagnostic performance of each gene. Evaluation metrics included AUC values, accuracy, sensitivity, and specificity. Diagnostic accuracy was categorized as follows: AUC between 0.5–0.7 was considered low accuracy, 0.7–0.9 as moderate accuracy, and greater than 0.9 as high accuracy.

### Assessment of immune cell infiltration

To evaluate the cellular composition of the immune microenvironment in OM samples, this study employed the CIBERSORT algorithm (43) to analyze immune cell infiltration based on gene expression profiles. Using the LM22 signature matrix, this algorithm deconvolutes the relative proportions of 22 immune cell subtypes within each sample, enabling a systematic assessment of immune cell infiltration patterns. Subsequently, the R package "ggplot2" was utilized to visualize the immune cell infiltration proportions, and box plots were generated to compare the infiltration levels of each immune cell subtype between OM patients and healthy controls.

### Intersection genes from network pharmacology and bioinformatics

To identify the intersection genes relevant to UA and OM, we utilized the R package ggvenn to intersect the disease-predicted genes from network pharmacology with the genes identified through ML for OM. These intersection genes were selected for subsequent analysis.

### Molecular docking

In the three-dimensional structure of proteins, ligand-binding sites and catalytic active sites are often located in surface structural pockets or internal cavities. In this study, we used the crystal structures in the Protein Data Bank (PDB) that already contain natural ligands as references, and their binding cavities were used as the preset sites for molecular docking. To ensure the accuracy of the binding site definition, the simulated protein structures were spatially superimposed with their original PDB structures in PyMOL (44), ensuring that the identified sites are consistent with the locations of the natural ligands. This validation step is crucial for maintaining the structural reliability and functional relevance of molecular docking, and it also helps enhance the understanding of protein spatial structure and potential binding domains. Molecular docking is a key computational technique for predicting the binding affinity between small molecules and proteins, capable of generating multiple ligand-protein complex conformations and ranking them based on binding free energy. By evaluating the binding modes and types of interactions between ligands and target proteins, this study aims to screen for potential candidate drugs and further explore the molecular recognition mechanisms between ligands and receptors.

### Molecular dynamics simulation

A 100 ns MD simulation of the complex was performed using Gromacs 2022 (45). The protein was described using the CHARMM36 force field parameters (46), while the ligand topology was generated with the GAFF2 force field (47). Periodic boundary conditions were applied, and the protein-ligand complex was solvated in a cubic box filled with TIP3P water molecules, maintaining a minimum distance of 1.2 nm between the solute and the box boundary. Electrostatic interactions were treated using the Particle Mesh Ewald (PME) method, and the Verlet cut-off scheme was employed. The system underwent energy minimization followed by equilibration under NVT (constant number of particles, volume, and temperature) and NPT (constant number of particles, pressure, and temperature) ensembles for 100 ps each, with a coupling constant of 0.1 ps. Both van der Waals and Coulomb interactions were calculated with a cut-off of 1.0 nm. Finally, a production MD simulation was carried out for 100 ns under constant temperature (310 K) and pressure (1 bar).

To evaluate the binding energy of the protein-ligand complex, the Molecular Mechanics/Poisson-Boltzmann Surface Area (MMPBSA) method was employed, and the binding affinity was calculated using the g_mmpbsa tool in GROMACS. The root-mean-square deviation (RMSD) was calculated to determine the equilibration time and assess the stability of each complex; lower RMSD values indicate better stability. The root- mean-square fluctuation (RMSF) was used to detect regions of high fluctuation, with higher RMSF values suggesting lower stability of the protein-ligand complex. The radius of gyration (Rg) was employed to evaluate the compactness of the protein structure; smaller Rg values indicate a more compact system. Finally, the solvent-accessible surface area (SASA) was calculated to assess hydrophobic interactions, with lower SASA values indicating stronger hydrophobic interactions and higher stability of the protein-ligand complex (48).

### Antibacterial experiments

#### Strain isolation

From September 1, 2023, to December 31, 2024, a total of 18 clinical strains were isolated from inpatients at the Hospital of Integrated Traditional Chinese and Western Medicine in Shenzhen. These strains were identified using the VITEK^®^ 2 Automated Compact System for Microbial Identification and Antimicrobial Susceptibility Testing (VITEK-2 system, bioMérieux, Inc., France).

#### Antibacterial agents

UA: Analytical standard, batch number C15297808, purity 98%, 1 g per vial, purchased from Shanghai Macklin Biochemical Technology Co., Ltd. Penicillin Sodium: Batch number J4012108, purity 98%, 80 units per vial, purchased from North China Pharmaceutical Co., Ltd. Mueller-Hinton Broth (MHB): Purchased from Beijing AoBoXing Biotechnology Co., Ltd.

#### Preparation of bacterial suspensions

Prior to experimentation, the cryopreserved strains were revived and transferred to plates for isolation and cultivation. One to two colonies with consistent morphology from a single plate were selected and inoculated into MHB, followed by incubation at 37°C for 10–12 hours to prepare the seed cultures. RPMI 1640 liquid medium was used to adjust the bacterial concentration, resulting in a bacterial suspension with a concentration of 2–6×10⁸ CFU/ml, which was designated as the stock bacterial suspension. For experimental use, the stock suspension was diluted 100 times with RPMI 1640 broth to achieve a final inoculum concentration of 2–6×10⁶ CFU/ml.

The drug solutions were prepared as follows: UA and penicillin were dissolved in DMSO and normal saline respectively to prepare mother solutions with a 256 µg/mL concentration. According to the microdilution method recommended by CLSI, the concentration gradient of the drugs was prepared as follows: The initial concentration of UA was 2048 µg/mL. A volume of 256 µg/mL drug solution was added to the first well of a 96- well plate, and then serial dilutions were performed with MHB. The concentration of the drug in the eighth well was 1 µg/mL, resulting in a concentration gradient of 1–128 µg/mL. The penicillin mother solution was also processed by serial dilution, yielding a final concentration gradient of 1–256 µg/mL.

#### Determination of MICs of UA and penicillin alone

The minimum inhibitory concentration (MIC) was determined according to the CLSI 2022 guideline, using the broth microdilution method. The specific steps are as follows: A 24-h MHB culture was prepared as the inoculum. The turbidity of the inoculum was adjusted to 0.5 McFarland units (approximately equivalent to 1.5×10⁸ CFU/mL) using normal saline, and then diluted to a final inoculum concentration of 1.5×10⁶ CFU/mL.

The susceptibility tests of UA and penicillin were performed in 96-well V-bottom plates. The drug mother solutions were serially diluted with MHB. The concentration of the drug in the first well was 256 µg/mL, and that in the eighth well was 1 µg/mL (for UA) or the corresponding gradient (for penicillin). Each well was inoculated with 100 µL of the diluted bacterial suspension, resulting in a final inoculum of approximately 1×10⁵ CFU per well. The 11th well served as the growth control (bacterial suspension only), and the 12th well as the blank control (broth only). The microtiter plate was covered and incubated at 37°C for 18–24 h. The MIC was defined as the lowest drug concentration at which no visible bacterial growth was observed to the naked eye.

#### Combination of UA and penicillin

The combined antibacterial effect of UA and penicillin was evaluated using the checkerboard microdilution method. A total of eight dilutions were prepared for each drug, with the highest concentration set at 4 × MIC, followed by two-fold serial dilutions (1, 1/2, 1/4, 1/8, 1/16, 1/32, 1/64, 1/128, and 1/256). The diluted drugs were combined in a 96-well plate along the horizontal and vertical axes, respectively. The bacterial inoculum was adjusted to 1×10⁵ CFU/mL. After incubating *S. aureus* at 37 °C for 18–24 hours, bacterial growth was assessed.

The fractional inhibitory concentration index (FICI) was used to evaluate the combined antibacterial effect. The FICI was calculated as follows:

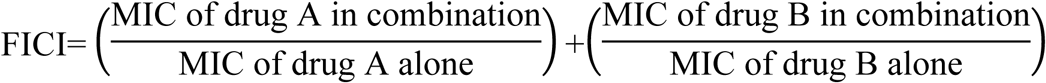

The results were interpreted according to the following criteria: Synergistic effect: FICI ≤ 0.5; Additive effect: 0.5 < FICI ≤ 1; Indifferent effect: 1 < FICI ≤ 2; Antagonistic effect: FICI > 2 (49).

All independent experiments were performed in triplicate.

#### Time-growth curve

Based on the monotherapy MIC results, the following experimental groups were established: UA (1/2 × MIC), Penicillin (1/4 × MIC), Penicillin (1/8 × MIC), UA + Penicillin (1/4 × MIC + 1/4 × MIC), UA + Penicillin (1/4 × MIC + 1/8 × MIC), UA + Penicillin (1/16 × MIC + 1/8 × MIC), UA + Penicillin (1/16 × MIC + 1/4 × MIC), UA (1 × MIC), Penicillin (1 × MIC), and a blank control. Each group was co-cultured with MRSA. MHB containing a final bacterial concentration of 1 × 10⁵ CFU/mL served as the control. Cultures were incubated at 37 °C with constant shaking. Bacterial growth was monitored by measuring the optical density at 600 nm (OD₆₀₀) at 0, 2, 4, 6, 8, 12, 14, and 24 hours. At each time point, 200 μL of bacterial suspension was transferred to a 96-well microplate, and the OD was recorded using a microplate reader. All assays were performed in triplicate wells, and the entire experiment was independently repeated three times.

#### Ethical Statement

All clinical strains involved in this study were derived from archived samples of inpatients at Shenzhen Hospital of Integrated Traditional Chinese and Western Medicine during the period from September 1, 2023, to December 31, 2024. The data were accessed for research purposes between January 2024 and December 2024. All patient identification information has been de-identified, and the researchers are unable to identify individual participant information. This study was approved by the Ethics Committee of Shenzhen Hospital of Integrated Traditional Chinese and Western Medicine (Approval No.: [KY-2023-082-02]).

### Statistical analysis

Statistical analyses were performed using SPSS software (version, 27.0) for analysis of antimicrobial susceptibility results. Intergroup comparisons were conducted using an independent *t*-test, with *P* < 0.05 considered statistically significant.

## Results

### Network pharmacology analysis

To explore the potential molecular targets of UA in the treatment of *S. aureus*-infected OM, network pharmacology analysis was performed. First, 75 UA-related targets (molecular formula: C[C@@H]1CC[C@@]2(CC[C@@]3(C(=CC[C@H]4[C@]3(CC[C@@ H]5[C@@]4(CCC@@HO)C)C)[C@@H]2[C@H]1C)C)C(=O)O) were screened using the SwissTargetPrediction database, and 267 related targets were identified from PharmMapper. After removing duplicates and taking the union of these two sets, 316 intersecting genes were obtained. Meanwhile, 2,626 genes associated with *S. aureus* infection and 1,529 genes related to OM were retrieved from the GeneCards, OMIM, and DisGeNET databases. A Venn diagram was constructed using a microbiology platform, leading to the identification of 57 common therapeutic targets (Fig 2A). To further visualize the regulatory effect of UA on *S. aureus*-infected OM, a PPI network was built based on the aforementioned intersecting targets, which contained 57 nodes and 453 edges (Fig 2B).

**Fig 2.**
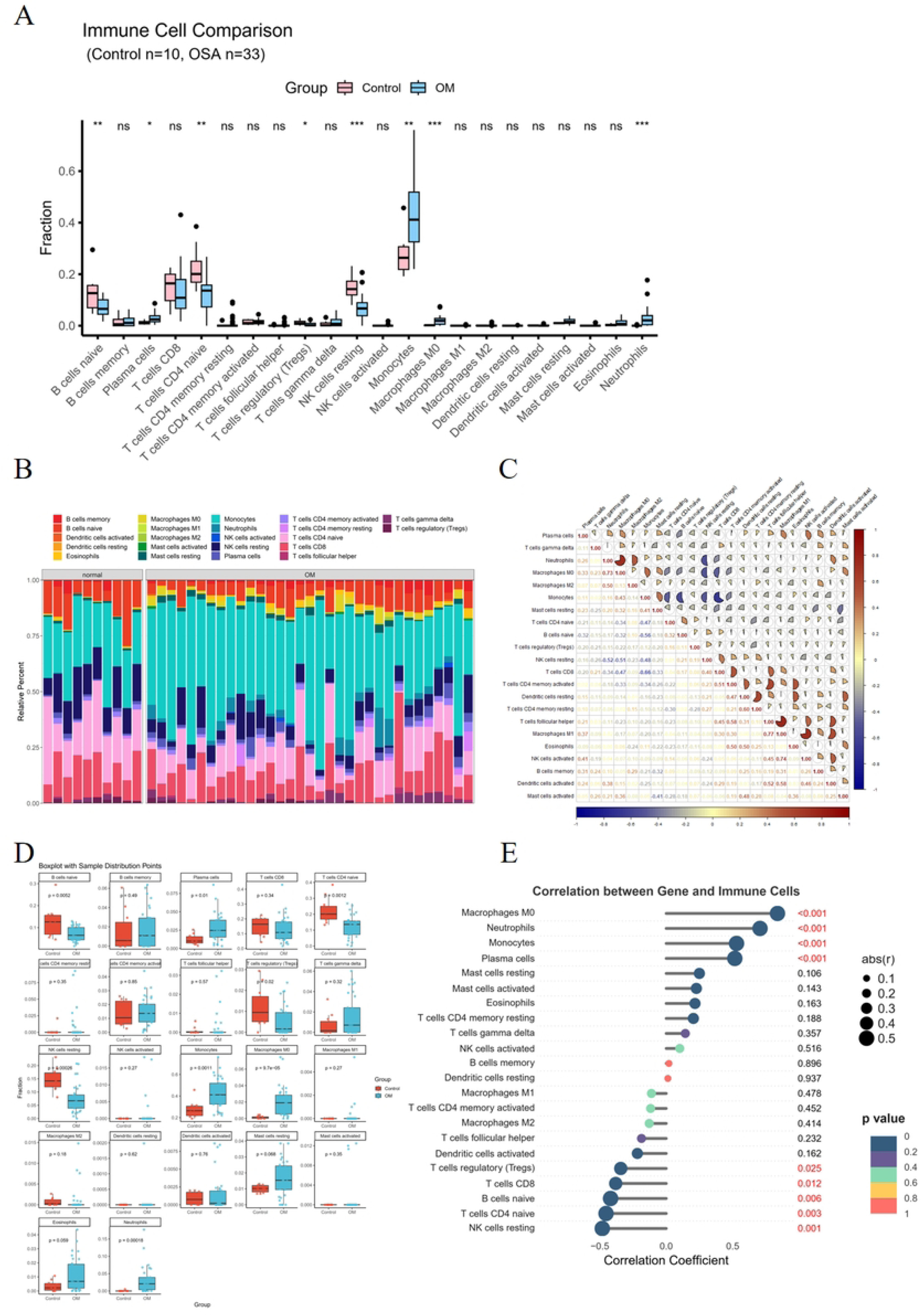
Target Screening and Functional Enrichment Analysis of Ursolic Acid for the Treatment of Staphylococcus aureus-Induced Osteomyelitis. (A) Identification of common targets of UA-OM-SAI. SAI: *S. aureus* infection. (B) PPI network of 57 common target genes, with a medium confidence interaction score of 0.400. PPI, protein-protein interaction. (C-E) The top 10 GO terms in BP, CC, and MF enrichment analysis, respectively. (F) The top 10 KEGG terms in the KEGG enrichment analysis.

To comprehensively and systematically elucidate the potential mechanism of UA in the treatment of OM, we performed a multi-level GO enrichment analysis of its therapeutic targets using the DAVID database, covering three categories: BP, CC, and MF. A total of 149 statistically significant GO terms were identified, including 1082 BP terms, 41 CC terms, and 87 MF terms. The top 10 enriched terms in BP, CC, and MF with the highest number of genes are presented in bar graphs (Fig 2C-E). The results indicate that the targets of UA in the treatment of OM were primarily enriched in BP such as regulation of inflammatory response, regulation of body fluid levels, positive regulation of cytokine production, and wound healing. Significant enrichment was also observed in CC including membrane raft, membrane microdomain, caveola, and vesicle lumen. Furthermore, these targets exhibited MF such as non-membrane spanning protein tyrosine kinase activity, transmembrane receptor protein tyrosine kinase activity, protein tyrosine kinase activity, and transmembrane receptor protein kinase activity.

To systematically investigate the intrinsic mechanisms underlying the anti-OM effects of UA, we conducted KEGG pathway analysis to predict signaling pathways associated with its potential therapeutic targets. The results revealed 151 statistically significant pathways. The top 10 pathways with the highest number of associated genes are displayed in a bar graph (Fig 2F), among which the Relaxin signaling pathway, Coronavirus disease – COVID-19, and PD-L1 expression and PD-1 checkpoint pathway in cancer were the most enriched. Based on the enrichment analysis of core targets and pathways, we focused on neutrophil extracellular trap (NET) formation and constructed a schematic diagram illustrating the mechanism by which UA treats OM by targeting this pathway (Fig 3). These findings suggest that UA exerts anti-OM effects through multiple pathways, with NET formation playing a critical role in its therapeutic action.

**Fig 3.**
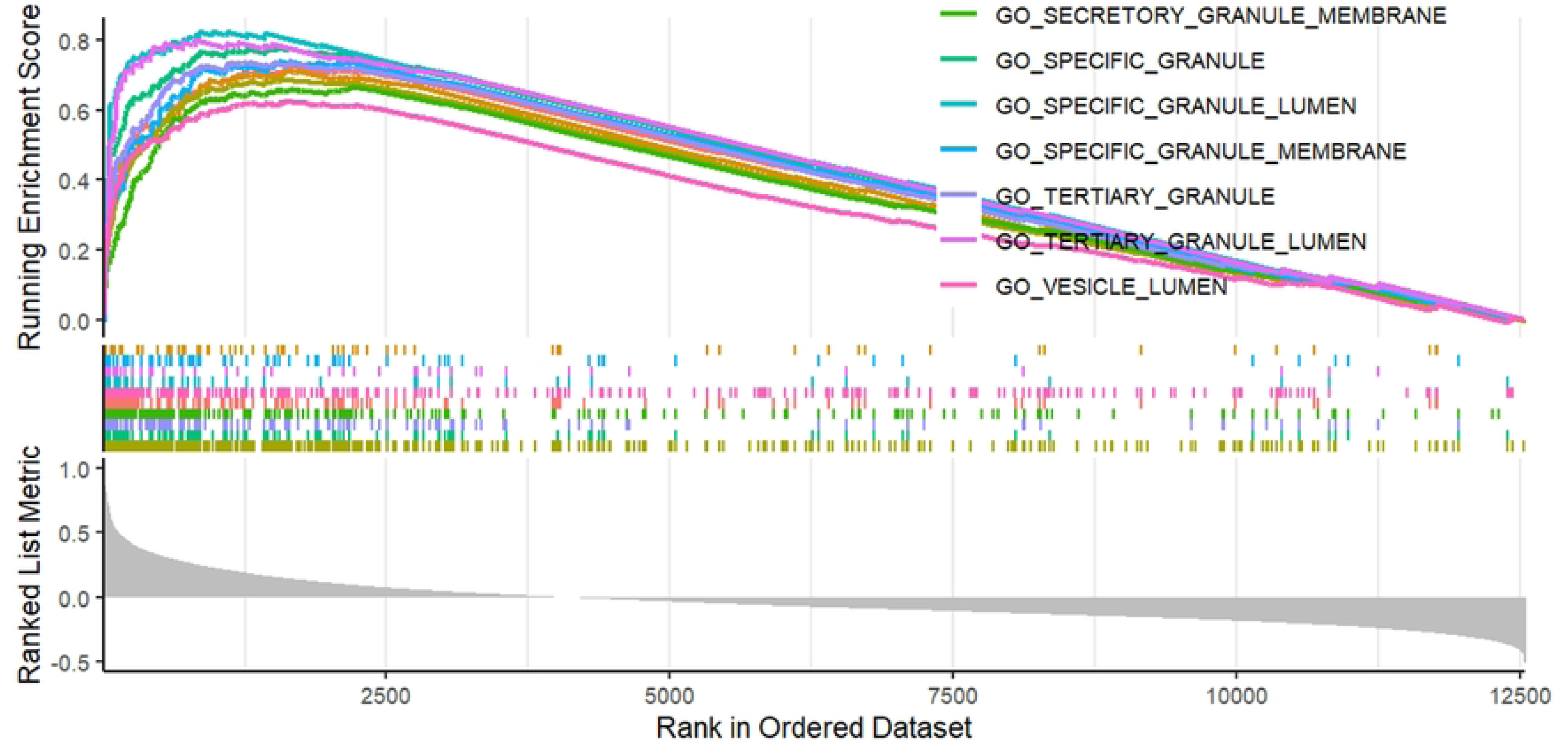
**Schematic illustration of the NET formation pathway.**

### WGCNA

WGCNA was employed to identify gene co-expression modules associated with patients with OM caused by *S. aureus* infection. After constructing a scale-free network using a soft-thresholding power of β = 4 (Fig 4A), a dynamic tree-cutting algorithm identified a total of 7 gene modules (gray, yellow, green, teal, red, blue, and brown modules) (Fig 4B), and their interaction relationships are illustrated in Fig 4C. A key finding revealed that among all modules, the blue module exhibited a significant correlation (Fig 4D). Further analysis demonstrated that this blue module was significantly correlated with GS (correlation coefficient = 0.60, *P* = 2.0×10⁻⁵) (Fig 4E); thus, it was identified as the key functional module, containing a total of 447 module genes.

**Fig 4.**
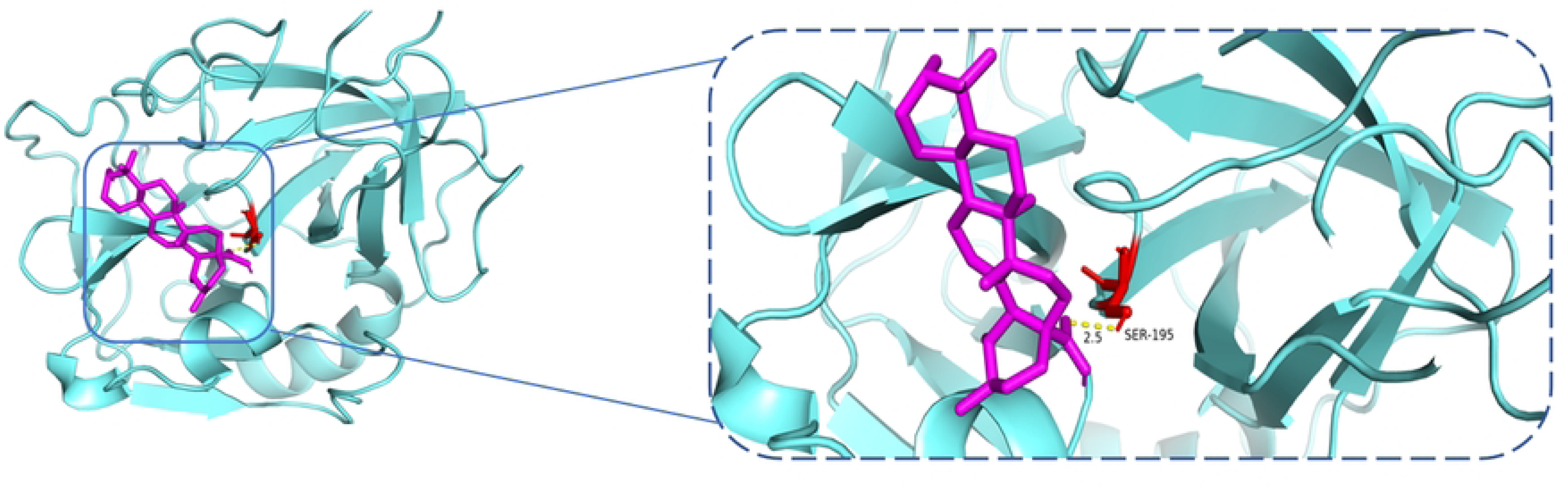
Identification of OM-related modules. (A) Analysis of soft-thresholding powers for achieving scale-free topology fit indices in the network. (B) Gene clustering dendrogram. (C) Heatmap depicting the relationships among different modules. (D) Heatmap illustrating module– trait associations. (E) Scatter plot showing the correlation between gene membership in the blue module and gene significance.

### Identification of DEGs

After normalizing the expression profiles of the GPL96 platform in the GSE16129 dataset (Fig 5A), 863 DEGs between the treated control group and the disease group were identified via volcano plot analysis (Fig 5B). Fig 5C presents the heatmap clustering results of the top 30 significantly differentially expressed DEGs. Furthermore, through integrated analysis of the DEGs and the most relevant module from WGCNA, we finally identified 64 core differential genes closely associated with OM.

**Fig 5.**
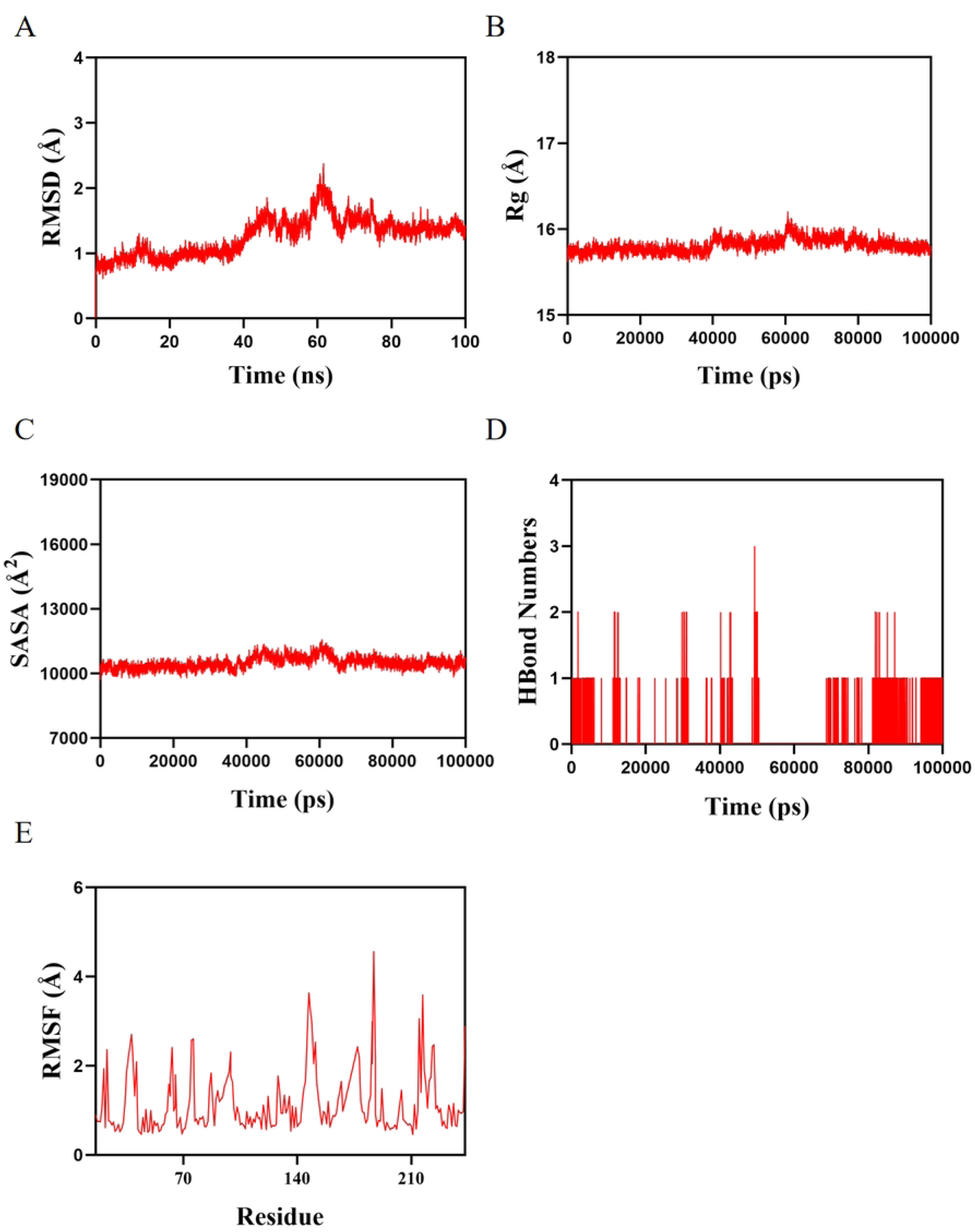
Identification of DEGs in OM. (A) PCA plot. (B) Volcano plot of DEGs in OM. (C) Heatmap of the top 30 DEGs in OM.

### Enrichment analysis of the intersection between DEGs and genes from the most relevant module

Functional annotation analysis was performed on the intersecting genes to clarify their potential biological functions. As shown in Fig 6A, in terms of BP, the DEGs were mainly involved in defense response to bacteria, humoral immune response, and antimicrobial humoral response. CC analysis revealed (Fig 6B) that these intersecting genes were associated with secretory granule lumen, cytoplasmic vesicle lumen, and specific granule. At the MF level (Fig 6C), the DEGs were significantly enriched in functions such as serine-type endopeptidase activity, serine-type peptidase activity, serine hydrolase activity, and glycosaminoglycan binding. In addition, pathway enrichment analysis (Fig 6D) indicated that the key intersecting genes were closely related to pathways, including Immune system and Cancer: overview. Further disease association analysis showed (Fig 6E) that these genes were significantly associated with diseases such as Coronavirus infectious disease, viral infectious disease, chronic obstructive pulmonary disease, and obstructive lung disease (Fig 6F and G).

**Fig 6.**
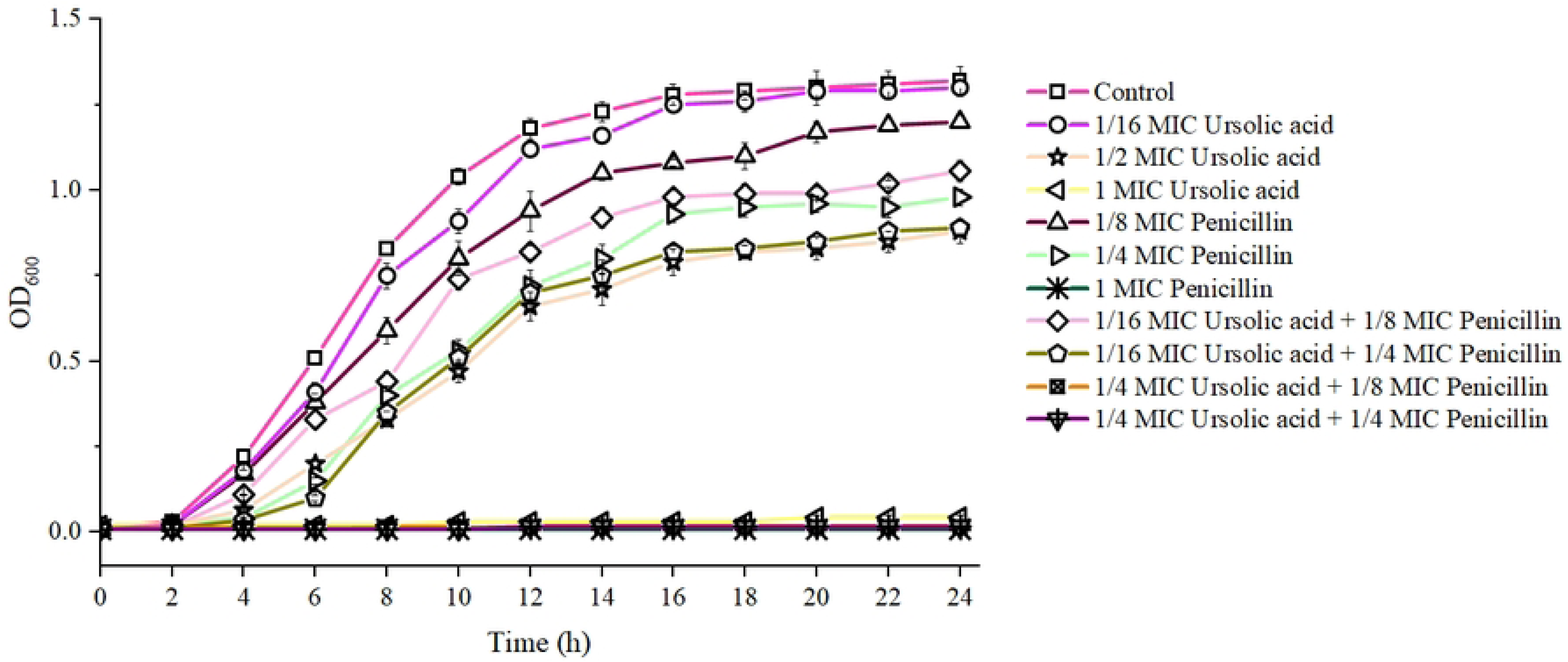
Top 10 functional enrichment terms of DEGs in OM. (A) Intersection of differentially expressed genes and the most relevant WGCNA module. (B) GO enrichment analysis in the BP category. (C) GO enrichment analysis in the CC category. (D) GO enrichment analysis in the MF category. (E) KEGG pathway enrichment analysis. (F) Gene-pathway association chord diagram. (G) DO enrichment analysis.

### PPI network and hub gene analysis

Genes typically do not act in isolation, as the proteins they encode can interact with one another. To explore the interaction relationships between proteins, a PPI network was constructed using the STRING database based on the key module genes identified by WGCNA, with a confidence score threshold set at 0.4. This network consisted of 46 nodes and 268 edges (Fig 7A).

**Fig 7.**
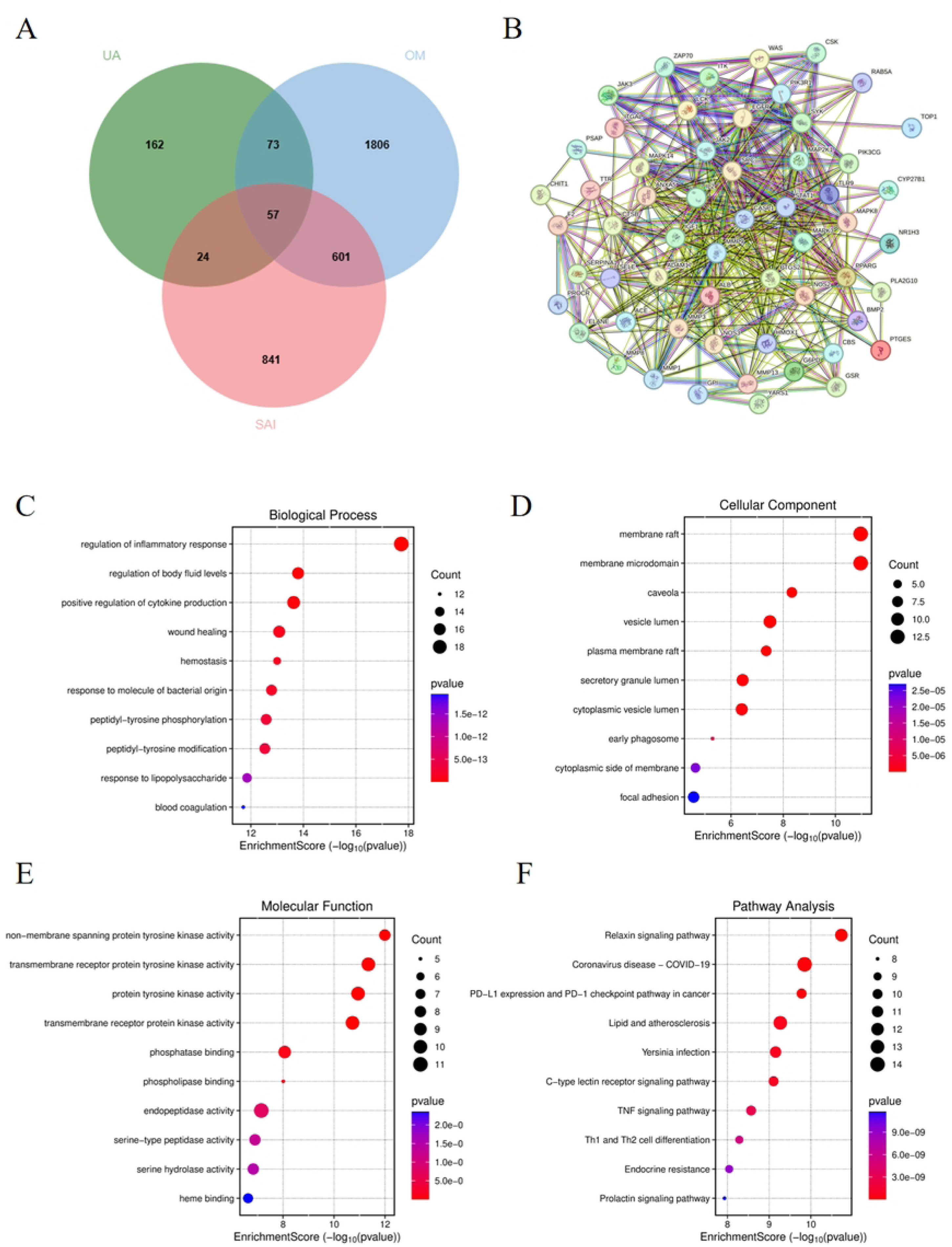
PPI network and hub gene analysis. (A) PPI network of DEGs in OM. The edge size corresponds to the degree of interaction. (B) The first functional module identified within the PPI network. (C) Identification of 19 common hub genes using four algorithms from the cytoHubba plugin.

The MCODE plugin was used to identify the most densely connected module within the network, which contained 20 nodes and 162 edges (Fig 7B). Additionally, four algorithms from the cytoHubba plugin were employed to screen the top 20 hub genes. Through comprehensive bioinformatics analysis, 19 common hub genes were selected, including *PRTN3*, *RNASE3*, D*EFA4*, *MS4A3*, *RNASE2*, *LTF*, *AZU1*, *MMP9*, *ELANE*, *CAMP*, *CEACAM8*, *MMP8*, *MPO*, *RETN*, and *CTSG* (Fig 7C).

### Identification and validation of key genes

Two ML algorithms were employed. The LASSO regression algorithm screened three potential diagnostic biomarkers (Fig 8A and B), while the RF algorithm identified ten diagnostic genes (Fig 8C and D). Venn diagram analysis ultimately determined three key genes—LTF, S100A12, and ELANE—as core diagnostic biomarkers (Fig 8E). An artificial neural network (ANN) diagnostic model was constructed based on gene weights (Fig 8F). ROC curve validation demonstrated excellent diagnostic performance for all three genes in OM (AUC values: LTF: 0.97, S100A12: 0.906, ELANE: 0.976) (Fig 9A). Further analysis revealed that all key genes were significantly differentially expressed in OM patients (Fig 9B) and exhibited strong diagnostic value for OM (Fig 9C and D).

**Fig 8.**
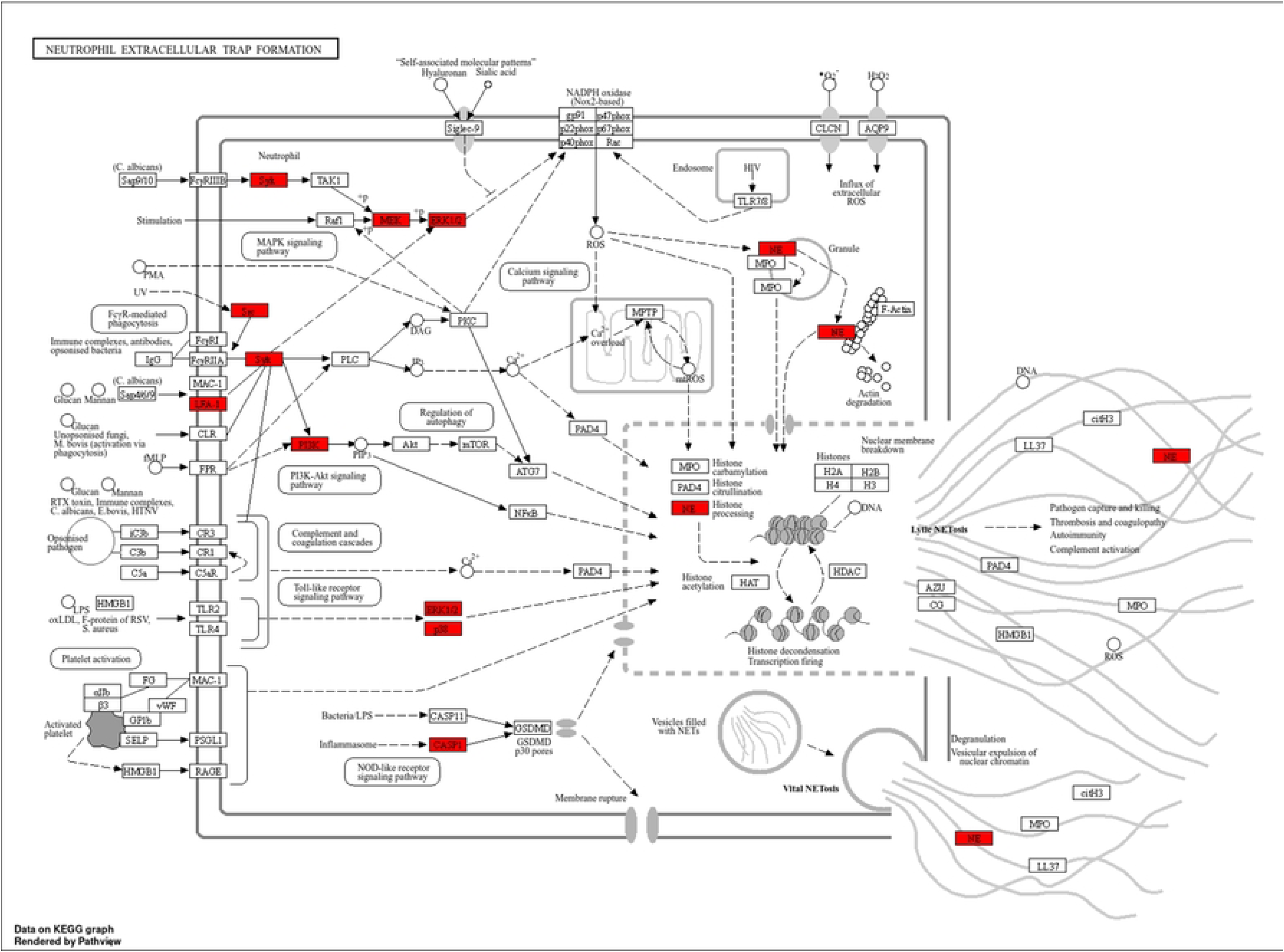
Identification of key genes in OM. (A-B) Three key genes identified from hub genes using LASSO regression. (C-D) Ten key genes identified from hub genes via the RF algorithm. (E) Overlap analysis revealing three consensus key genes: ELANE, LTF, and S100A12. (F) ANN model constructed for the key genes.

**Fig 9.**
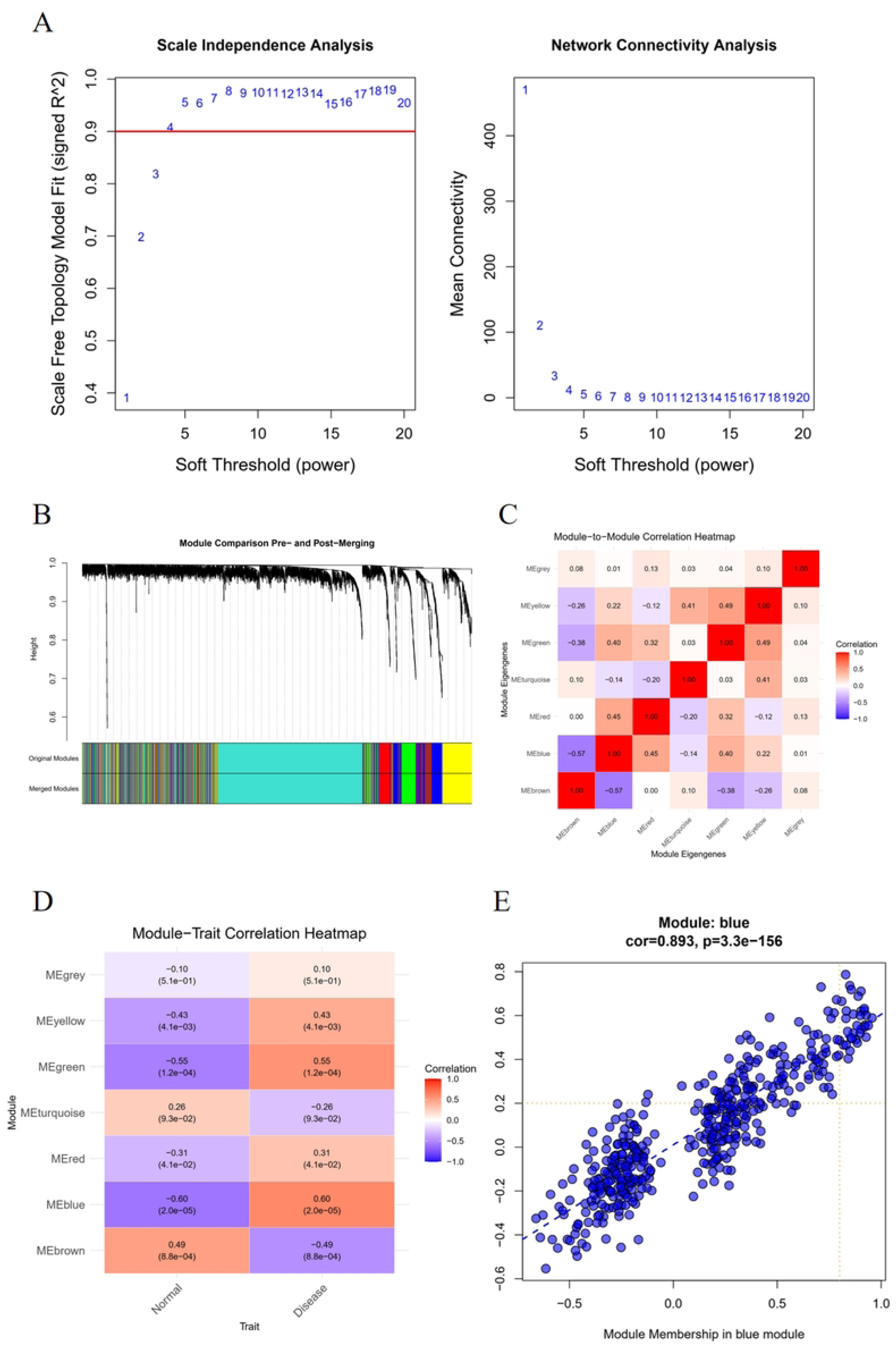
Validation of key genes in OM. (A) ROC analysis of key genes in the ANN model. (B) Significant expression of key genes in OM. (C) ROC curves of key genes in OM. (D) Forest plot of key genes in OM.

ELANE is a protease predominantly secreted by neutrophils that plays a critical regulatory role in inflammation and immune responses. As a member of the serine protease family, its primary function involves the degradation of extracellular matrix proteins, thereby facilitating essential BP such as tissue remodeling, leukocyte migration, and cell signaling during inflammation. Additionally, ELANE contributes to the modulation of inflammatory responses through the regulation of inflammatory mediator release, cleavage of pro-inflammatory cytokines, and liberation of soluble complement receptors that inhibit complement activation. These multifaceted functions underscore its importance in the fine-tuning and resolution of inflammatory processes.

LTF, an iron-binding protein belonging to the transferrin family, exerts multiple biological effects in humans, including antibacterial, antioxidant, antitumor, antiviral, immunomodulatory, and anti-aging activities (50, 51). Beyond regulating lipid metabolism and reducing fat accumulation, LTF promotes osteogenic differentiation and suppresses oxidative stress (OS) (52, 53).

S100A12, a newer member of the S100 calcium-binding protein family, is located within the gene cluster on chromosome 1q21 along with other S100A family members. In humans, S100A12 is mainly secreted by neutrophils and monocytes, though it is also expressed in tissue cells such as mucosal epithelial cells and keratinocytes (note: rodents do not express S100A12). Its upregulation is primarily driven by cellular stress, which induces the release of this protein from granulocytes and other myeloid cells(54).

### Immune infiltration analysis

Compared with the healthy control group, OM samples in the disease group exhibited a characteristic immune profile, with decreased proportions of naive B cells, plasma cells, naive CD4⁺ T cells, regulatory T cells (Tregs), and resting natural killer (NK) cells, while showing increased proportions of M0 macrophages, monocytes, and neutrophils (Fig 10A). Correlation analysis revealed that the key gene ELANE was significantly negatively correlated with naive B cells, CD8⁺ T cells, naive CD4⁺ T cells, Tregs, and resting NK cells, but positively correlated with plasma cells, monocytes, and M0 macrophages (Fig 10B). Subsequently, correlation heatmap analysis between immune cells revealed that in OM samples, T cells follicular helper showed a positive correlation with Macrophages M1 (r = 0.77), while T cells CD8 demonstrated a negative correlation with Monocytes (r = -0.66) (Fig 10C). These findings indicate distinct immune patterns in patients compared to controls, as well as interactions between various types of immune cells. In addition, correlation analysis revealed that ELANE showed significant correlations with Macrophages M0, Neutrophils, Monocytes, Plasma cells, T cells regulatory (Tregs), T cells CD8, B cells naive, T cells CD4 naive, and NK cells resting (Fig 10D and E).

**Fig 10.**
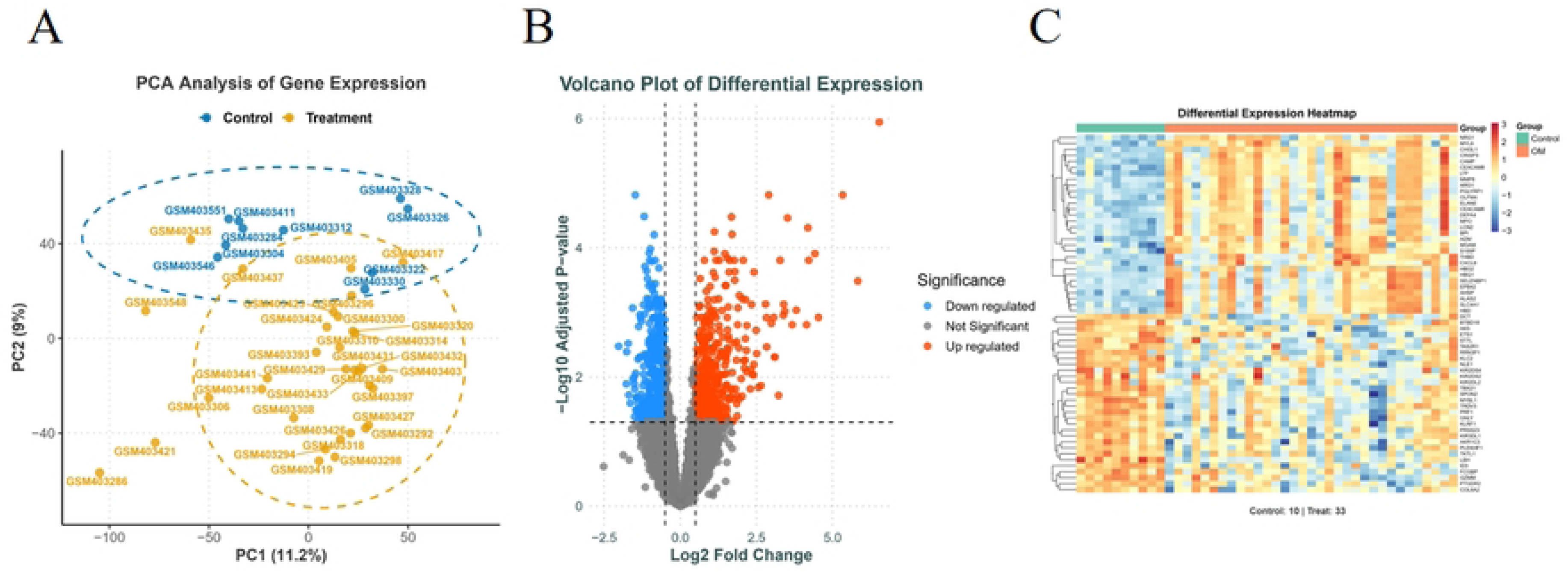
Immune infiltration. (A) Box plot of immune cell infiltration levels. (B) Stacked bar chart showing the composition of immune cell infiltration. (C) Heatmap depicting correlations between gene expression and immune cell infiltration. (D) Expression of immune cells in OM patients. (E) Relationship between ELANE and immune cells in OM patients. \**P* < 0.05, \*\**P* < 0.01, and \*\*\**P* < 0.001

UA can significantly inhibit the production of proinflammatory cytokines, such as tumor necrosis factor-α (TNF-α), IL-1β, and IL-6. For instance, in lipopolysaccharide (LPS)-induced RAW 264.7 macrophages, UA significantly reduced the mRNA expression levels of TNF-α and IL-6. Additionally, in LPS-activated macrophages, UA inhibited the activation of the nuclear factor-κB (NF-κB) signaling pathway, thereby reducing the release of inflammatory cytokines.

UA can regulate the polarization state of macrophages, promoting their transition from the proinflammatory M1 phenotype to the anti- inflammatory M2 phenotype. *In vitro* experiments have confirmed that UA can inhibit the polarization of M1 macrophages while promoting the polarization of M2 macrophages, thereby exerting anti-inflammatory effects. For example, in a diabetic wound model, UA-treated nanofibrous dressings significantly inhibited LPS-induced M1 macrophage polarization, restored the M2 phenotype, and accelerated the resolution of inflammation.

UA exerts anti-inflammatory effects by inhibiting the Toll-like receptor 4 (TLR4)/myeloid differentiation factor 88 (MyD88) signaling pathway. Furthermore, UA can reduce the production of inflammatory mediators by inhibiting the activation of the NF-κB and mitogen-activated protein kinase (MAPK) signaling pathways. In a chronic bronchitis model, for instance, UA alleviated inflammation by inhibiting the nuclear translocation of the NF-κB p65 subunit and downregulating the expression of TNF-α, IL-1β, prostaglandin E2 (PGE2), and leukotriene B4 (LTB4).

UA can enhance the autophagic activity of macrophages by upregulating the expression of autophagy-related genes Atg5 and Atg16L1, thereby regulating macrophage function and mitigating inflammatory responses. In an atherosclerosis model, UA reduced inflammation by enhancing macrophage autophagic activity and inhibiting M1 macrophage polarization.

The regulatory effect of UA on macrophages also exhibits significant anti-inflammatory effects in various tissues. In the liver, for example, UA alleviates liver fibrosis by inhibiting the pyroptosis of Kupffer cells and the NOX2/NLRP3 inflammasome signaling pathway. In the skin, UA can promote wound healing and inhibit inflammatory responses by regulating the polarization state of skin macrophages and promoting the accumulation of M2 macrophages.

### Gene Set Enrichment Analysis (GSEA) enrichment analysis of ELANE

Ultimately, the gene ELANE was identified as the intersection through the integration of network pharmacology and bioinformatics analyses, and was selected for subsequent investigations. To further elucidate the mechanism of action of ELANE—a key factor in OM—a multi-level regulatory network was constructed. GSEA revealed that ELANE is primarily involved in three major signaling pathways: Fcγ receptor (FcγR)- mediated phagocytosis, leukocyte transendothelial migration, and glycolysis/gluconeogenesis (Fig 11).

**Fig 11.**
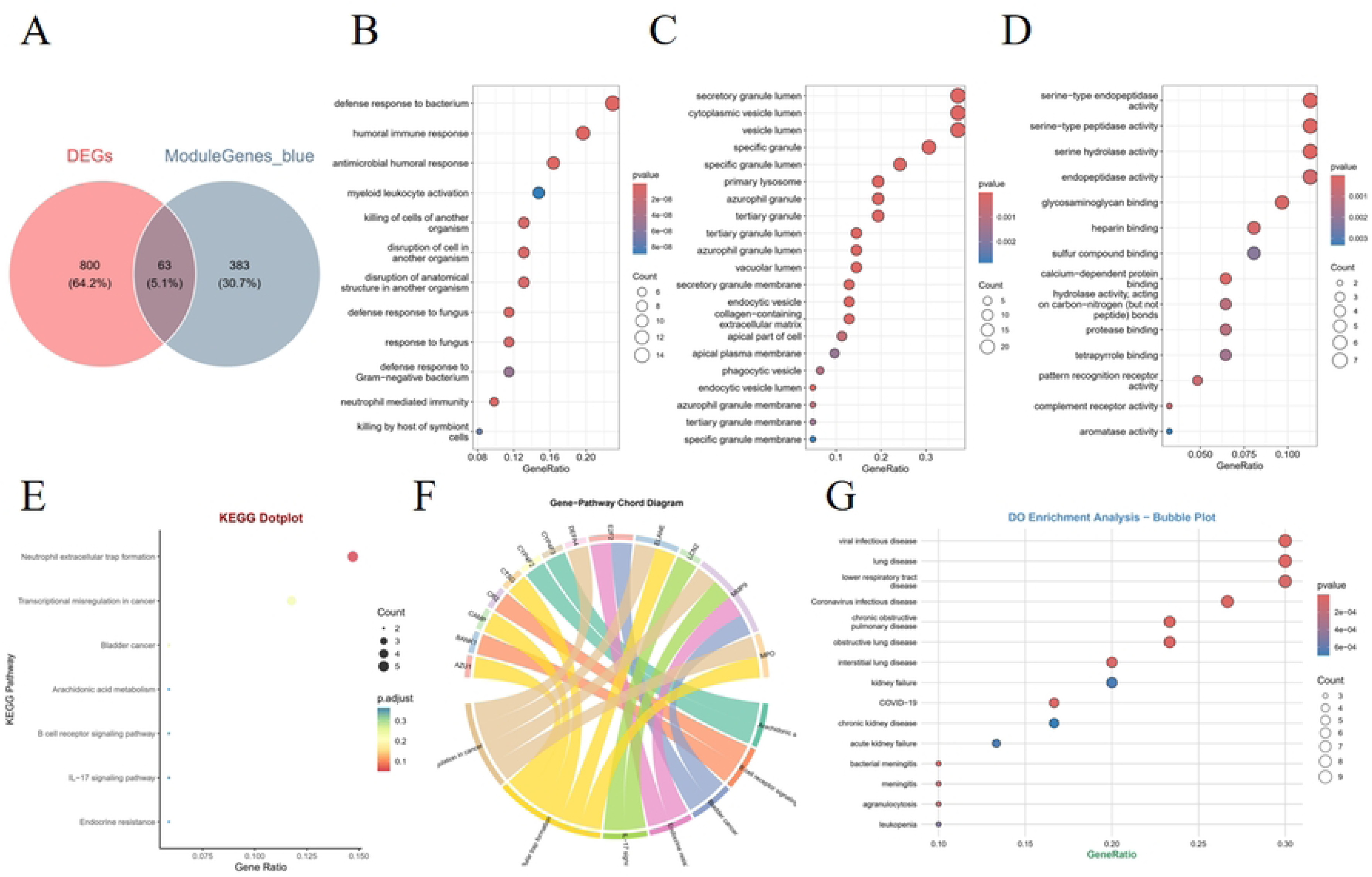
**GSEA of ELANE.**

### Molecular docking and molecular dynamics simulation

Molecular docking was performed using UA as the ligand and the key target ELANE as the receptor. The reliability of the docking procedure was first validated for the protein macromolecule. The RMSD values for ELANE were all ≤ 2.0 Å, confirming the reliability of the docking method. A higher CIE value indicates better docking activity. UA exhibited favorable binding affinity to the key target (Fig 12).

**Fig 12.**
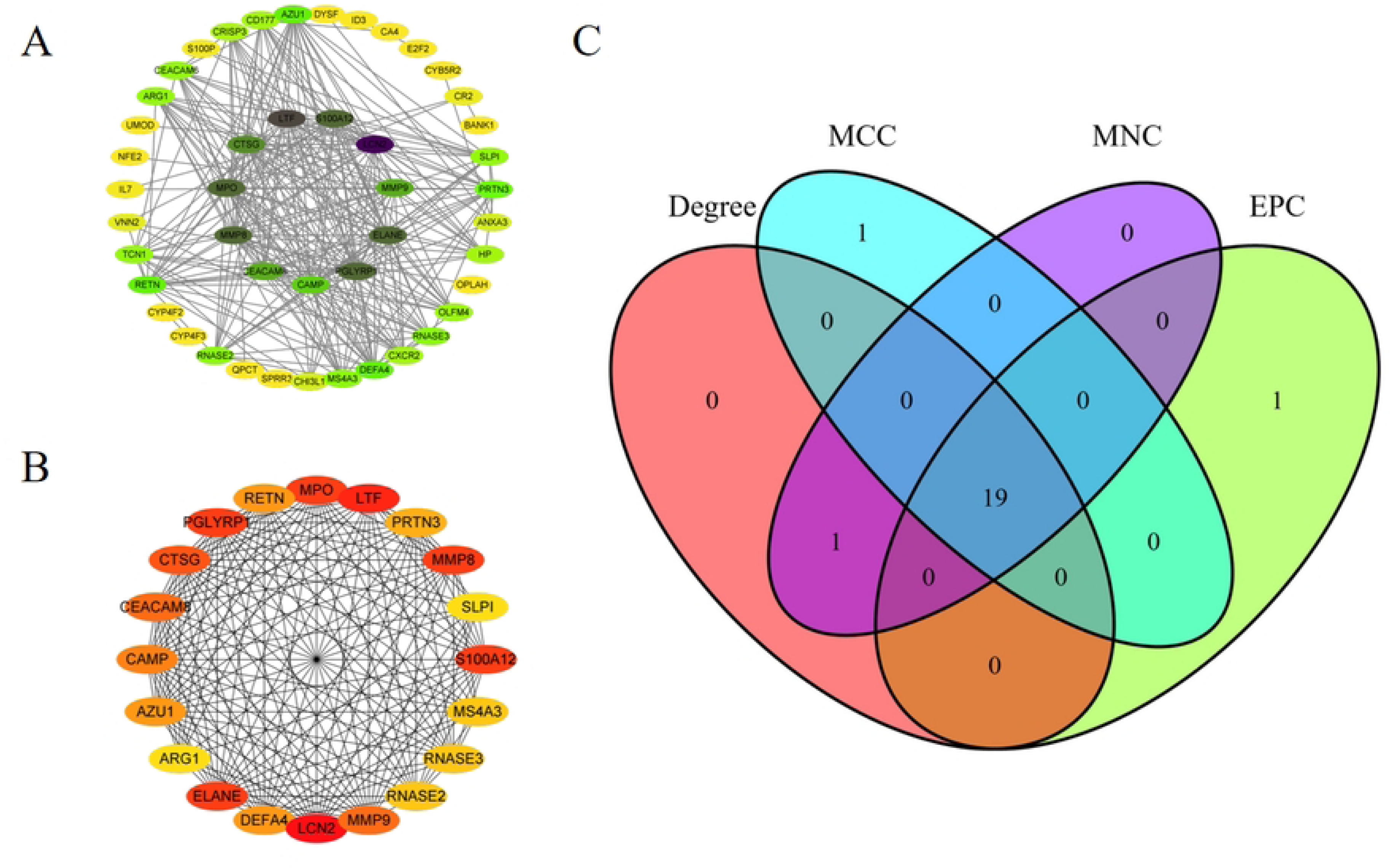
Nteraction diagram of UA with ELANE.

Smaller deviations indicate greater conformational stability. Thus, RMSD was used to evaluate the equilibrium of the simulation system. As shown in Fig 13A, the complex system reached equilibrium after 75 ns, eventually fluctuating around 1.4 Å, indicating high stability of the small molecule when bound to the target protein.

**Fig 13.**
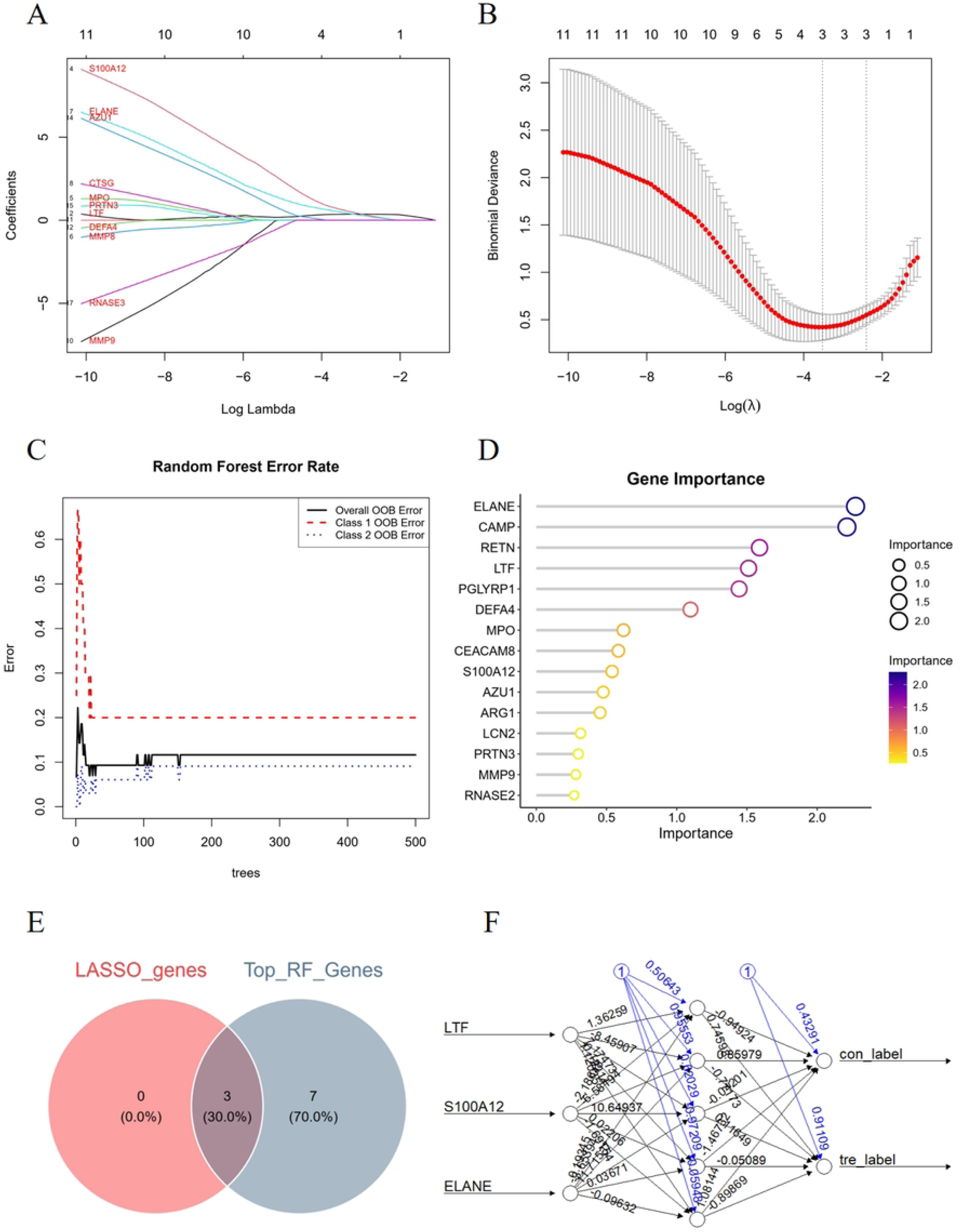
Molecular dynamics simulation analysis of the UA-ELANE complex. (A) Time-dependent RMSD of the protein–ligand complex. (B) Time-dependent Rg of the protein–ligand complex. (C) Time-dependent SASA of the protein–ligand complex. (D) Time-dependent number of hydrogen bonds in the protein–ligand complex. (E) Time-dependent RMSF of the backbone atoms of the protein–ligand complex.

The Rg describes changes in the overall structure and reflects the compactness of the protein. Greater variations in Rg suggest increased structural expansion. The complex system exhibited minor fluctuations during simulation, indicating conformational changes in the small molecule–target protein complex (Fig 13B). The SASA was used to evaluate the surface area of the protein. The SASA between the target protein and the small molecule was calculated during the simulation (Fig 13C). The results showed slight fluctuations followed by stabilization, demonstrating that ligand binding influences the micro-environment and leads to moderate changes in SASA.

Hydrogen bonds play a crucial role in ligand–protein binding. The number of hydrogen bonds between the small molecule and the target protein during dynamics is shown in Fig 13D. The complex system formed between 0 and 3 hydrogen bonds, with one bond observed in most cases, indicating favorable hydrogen bond interactions between the ligand and the target protein.

The RMSF reflects the flexibility of amino acid residues in the protein. As shown in Fig 13E, the complex exhibited relatively low RMSF values (mostly below 2.3 Å), suggesting low flexibility and high stability.In summary, the complex system demonstrated stable binding, supported by favorable hydrogen bond interactions, indicating strong binding between the small molecule and the target protein.

### MICs of UA and penicillin against MRSA

The MIC of UA against MRSA ranged from 16 to 64 μg/mL, while the MIC of penicillin against MRSA ranged from 32 to 512 μg/mL (Table 1).

**Table 1.** Synergistic effects of UA combined with penicillin against 18 MRSA isolates (Unit: μg/mL)

| MRSA | UA | Penicillin | FICI |
| --- | --- | --- | --- |

| Strain | MIC<sub>alone</sub> | MIC<sub>combination</sub> | MIC<sub>alone</sub> | MIC<sub>combination</sub> |  |
| --- | --- | --- | --- | --- | --- |
| 1 | 16 | 4 | 256 | 32 | 0.375 |
| 2 | 64 | 4 | 256 | 16 | 0.125 |
| 3 | 32 | 8 | 128 | 1 | 0.257 |
| 4 | 32 | 8 | 512 | 2 | 0.253 |
| 5 | 32 | 8 | 64 | 32 | 0.375 |
| 6 | 64 | 8 | 512 | 64 | 0.375 |
| 7 | 32 | 8 | 256 | 32 | 0.5 |
| 8 | 64 | 8 | 512 | 16 | 0.156 |
| 9 | 32 | 8 | 256 | 16 | 0.312 |
| 10 | 32 | 8 | 32 | 8 | 0.5 |
| 11 | 64 | 8 | 256 | 4 | 0.14 |
| 12 | 64 | 8 | 128 | 1 | 0.132 |
| 13 | 32 | 8 | 128 | 8 | 0.312 |
| 14 | 32 | 8 | 256 | 16 | 0.312 |
| 15 | 32 | 8 | 512 | 16 | 0.281 |
| 16 | 16 | 4 | 128 | 16 | 0.375 |
| 17 | 32 | 8 | 512 | 4 | 0.257 |
| 18 | 32 | 8 | 128 | 2 | 0.265 |

The FICI of the combination of UA and penicillin against MRSA was 0.125–0.5, indicating a synergistic effect between UA and penicillin in inhibiting MRSA. When the two drugs were used in combination, the optimal inhibitory concentration of UA was 75%–93.75% lower than the MIC of UA alone against MRSA, and the optimal inhibitory concentration of penicillin was 75%–99.3% lower than the MIC of penicillin alone against MRSA.

### Bactericidal curves of UA combined with penicillin against MRSA

The growth curve reflects the proliferation pattern of bacteria in a specific environment. The effects of UA at different concentrations on the growth curve of MRSA are shown in Fig 14. As observed, the control group exhibited a typical sigmoidal growth curve with distinct lag, exponential, and stationary phases, ultimately reaching a high bacterial density. In contrast, the addition of UA significantly inhibited the growth of MRSA in a concentration-dependent manner.

**Fig 14.**
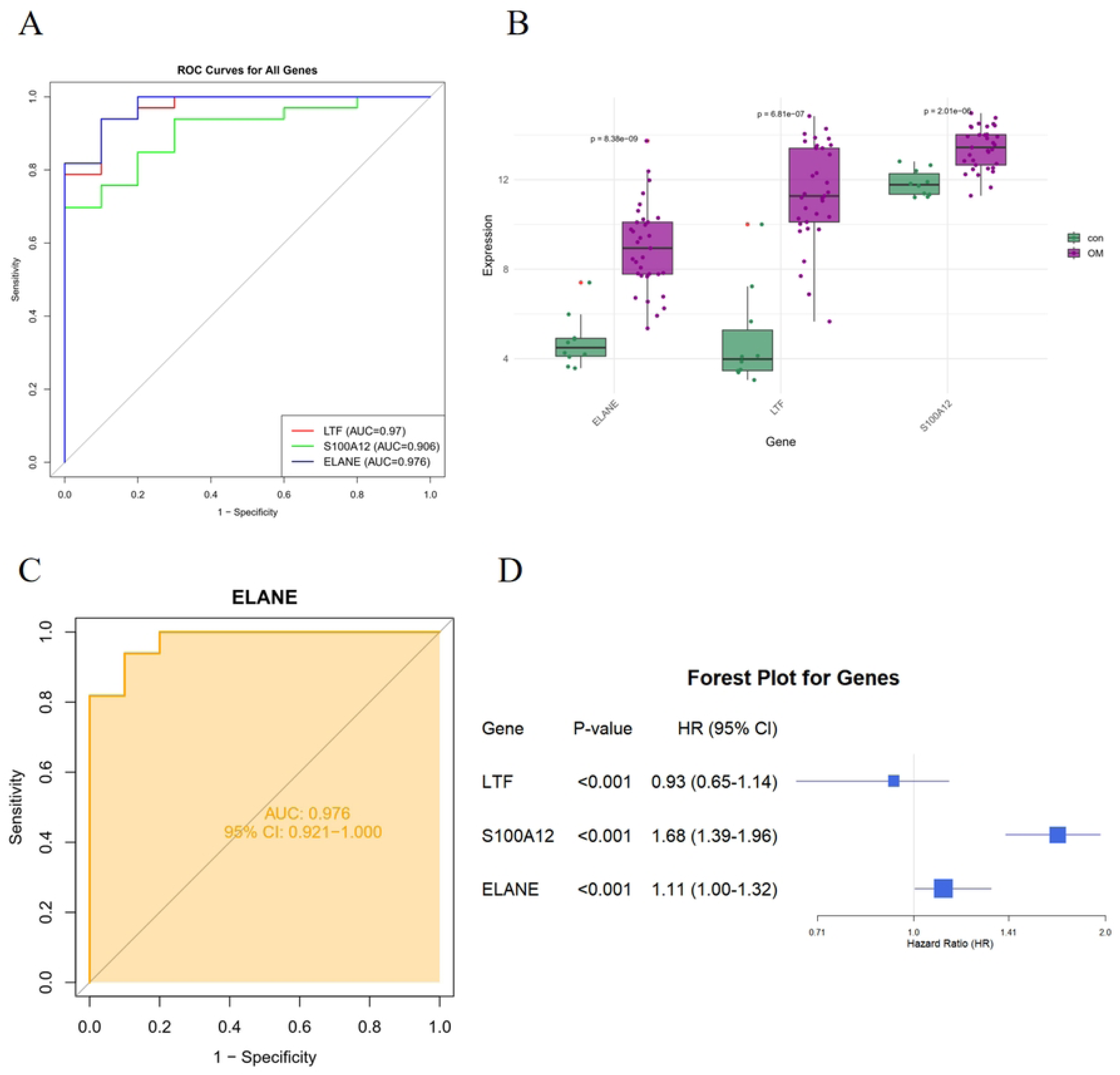
Bacterial growth curves at various concentrations.

At 1/16 × MIC, UA exerted a moderate inhibitory effect, though the bacteria were still able to enter the stationary phase, with a final bacterial density lower than that of the control. When the concentration of UA was increased to 1/2 × MIC, the inhibition during the exponential phase became more pronounced, resulting in a markedly reduced growth rate and a substantial decrease in final biomass during the stationary phase. Under treatment with 1 × MIC UA, the inhibitory effect was most robust, severely restricting bacterial proliferation throughout the entire growth cycle and almost completely suppressing the exponential phase.These results demonstrate that UA effectively disrupts the normal growth cycle of MRSA, significantly delaying its growth and reducing the final bacterial load.

## Discussion

Osteomyelitis(OM), particularly when induced by methicillin- resistant *Staphylococcus aureus* (MRSA), represents a formidable clinical challenge in orthopedics due to its complex pathogenesis and the escalating problem of antibiotic resistance(55). This study employed an integrated approach combining network pharmacology, bioinformatics, and *in vitro* experimental validation to systematically elucidate the multi-target mechanisms and synergistic antibacterial potential of ursolic acid (UA) against *S. aureus*-induced OM(56).

A pivotal finding was the identification and validation of three core genes—Neutrophil Elastase (*ELANE*), Lactoferrin (*LTF*), and *S100A12*— using machine learning algorithms (LASSO and Random Forest). These genes play central roles in the immune and inflammatory responses characteristic of OM. Functional enrichment analyses further revealed that the mechanisms of UA were significantly associated with key pathways, including neutrophil extracellular trap (NET) formation, inflammatory response, and immune regulation. Specifically, *ELANE*, a key protease secreted by neutrophils, is involved not only in extracellular matrix degradation and tissue remodeling but also in the fine-tuning of inflammatory responses through the regulation of inflammatory mediators and complement activation. Our molecular docking and dynamics simulations provided compelling structural biological evidence by demonstrating stable binding between UA and *ELANE*, supporting the novel mechanism of UA targeting NET formation.

Regarding antibacterial activity, UA exhibited moderate direct antibacterial effects against clinically isolated MRSA strains (MIC = 16–64 μg/mL). More importantly, the combination of UA with penicillin demonstrated a significant synergistic effect (FICI = 0.125–0.5), drastically reducing the effective inhibitory concentrations of both agents. This finding holds considerable clinical implications, suggesting UA’s potential as an antibiotic adjuvant to resensitize resistant bacteria like MRSA to conventional antibiotics, thereby offering a promising strategy to combat the global threat of antimicrobial resistance.

Immune infiltration analysis unveiled a distinct immune microenvironment in OM patients, characterized by a decreased proportion of naive lymphocyte subsets (e.g., naive B cells, CD4⁺ T cells, resting NK cells) and significant infiltration of myeloid cells (e.g., monocytes, M0 macrophages, neutrophils). Correlation analysis indicated a positive association between the core gene ELANE and these myeloid immune cells, further reinforcing the notion that UA may exert its therapeutic effects by modulating the function of neutrophils and other immune cells. Consistent with existing literature, which reports that UA can inhibit pro- inflammatory signaling pathways (e.g., NF-κB, MAPK), promote macrophage polarization from the M1 to the M2 phenotype, and enhance autophagy, our results collectively depict a coherent picture where UA combats OM through synergistic "immunomodulatory–anti- inflammatory–direct antibacterial" pathways(57, 58).

Nevertheless, this study has several limitations. First, the protein expression levels of core targets like *ELANE* require further validation using techniques such as Western blot or immunohistochemistry on tissue samples. Second, the diagnostic model constructed in this study necessitates validation in larger, independent cohorts before clinical application, due to the constraints of our initial sample size. Most crucially, the current conclusions are primarily based on bioinformatic predictions and *in vitro* experiments. The actual *in vivo* efficacy of UA and its therapeutic effects in animal models of OM remain to be determined, representing a critical gap that future research must address.

Looking forward, research on OM presents both challenges and opportunities. Clinically, sustained attention to multidisciplinary care guidelines and evolving antimicrobial susceptibility patterns is paramount(58, 59). From a basic research perspective, given that standard *in vitro* susceptibility testing fails to replicate the complexity of the *in vivo* infection microenvironment (e.g., biofilms, hypoxia), developing novel local and bone-targeted drug delivery systems will be a key future direction to overcome treatment barriers. Furthermore, evidence from animal models and human samples indicates that *S. aureus* can invade the osteocyte lacunar-canalicular network, and this spatial heterogeneity in colonization may contribute to treatment failure(60). Therefore, developing novel methodologies to investigate host-pathogen interactions in a spatially resolved manner will deepen our understanding of OM pathophysiology.

In conclusion, this study is the first to comprehensively reveal, from systems biology and experimental perspectives, that UA acts against OM through dual mechanisms—immunomodulation and direct antibacterial activity. It provides a theoretical foundation and experimental evidence for the potential use of UA as an adjunctive therapy for MRSA-related OM and highlights its role as a potential antibiotic sensitizer. Subsequent research should prioritize *in vivo* efficacy assessments and preclinical translational studies to advance UA as a candidate molecule for novel anti- osteomyelitic drugs.

## Conclusion

By integrating network pharmacology, bioinformatics analysis, and *in vitro* experimental validation, this study systematically elucidated the multi-target mechanism of action and synergistic antibacterial potential of UA in the treatment of OM caused by *S. aureus* infection. The findings revealed that UA may interfere with the pathological progression of OM by targeting key genes such as *ELANE*, *LTF*, and *S100A12*, and regulating critical pathways including NET formation, inflammatory responses, and immune modulation. Molecular docking and MD simulations further confirmed the stable binding ability of UA to core targets (particularly *ELANE*). In vitro antibacterial experiments demonstrated that UA exhibits moderate antibacterial activity against clinically isolated MRSA strains (MIC = 16–64 μg/mL) and exerts a significant synergistic effect when combined with penicillin (FICI = 0.125–0.5), substantially reducing the required concentration of antibiotics.

In conclusion, this study is the first to clarify that UA exerts anti-OM effects through dual mechanisms—immunomodulation and direct antibacterial activity—from the perspectives of systems biology and experimental validation. It provides a theoretical basis and experimental evidence for the potential use of UA as an adjuvant agent in the clinical treatment of MRSA infection-related OM. Future research should focus on further *in vivo* efficacy validation and clinical translation to advance UA as a candidate molecule for novel anti-OM drugs.

## Acknowledgments

Not applicable.

